# Genomic Evidence for Revising Management Units of European Anchovy: Integrating Evolutionary Lineages into Fisheries Assessment

**DOI:** 10.64898/2025.12.05.692497

**Authors:** Ane del Rio, Natalia Díaz-Arce, Leire Ibaibarriaga, María Santos-Mocoroa, Susana Garrido, Naiara Rodríguez-Ezpeleta

## Abstract

Effective fisheries management requires stock boundaries that accurately reflect underlying biological populations. In European anchovy (*Engraulis encrasicolus*), defining management units has been complicated by a complex evolutionary history and the coexistence of distinct ecotypes. Despite recent advances using high-resolution genomic markers, key uncertainties persist regarding population structure and connectivity, particularly between the Bay of Biscay and Atlantic Iberian Waters stocks and their links to neighboring northern and southern regions. Here, we analyze thousands of genetic markers from individuals spanning both assessed stocks and adjacent areas, including representatives of the two recognized ecotypes (marine and coastal). Our comprehensive population genomic analyses identify three major lineages, northern marine, southern marine, and coastal, shaped by historical processes and ecological differentiation. Notably, genetic divergence between ecotypes exceeds that observed among geographically distant populations within the same ecotype, highlighting the need to incorporate ecological as well as spatial drivers when delineating stocks. Our findings demonstrate that current management units do not capture the underlying biological structure of European anchovy, which could lead to local overexploitation due to inadequate Total Allowable Catch (TAC) settings. This study emphasizes the need to incorporate genetic data when defining biologically relevant management units, with the goal of improving stock assessments, safeguarding adaptive potential, and ensuring the long-term sustainability of the species.

## 1. INTRODUCTION

Properly defined stock boundaries are essential for effective fisheries assessment and management, as mismatches between biological populations and management units can lead to biased estimates of stock status and inappropriate harvest strategies (Kerr et al., 2017). This implies that stocks should represent natural populations composed of sexually interbreeding individuals sharing a common gene pool (Kerr et al., 2017). In practice, however, stocks are frequently delimited by administrative or political considerations and, for many species, there is evidence of misalignment between population and stock boundaries (e.g. Hutchinson (2008); Leone et al. (2019)). Such discrepancies can result in inaccurate interpretations of stock status, potentially leading to misguided management advice, which increases the risk of setting inappropriate Total Allowable Catch (TAC) limits, leading to local depletion or yield loss (Kerr et al., 2016; Reiss et al., 2009). Despite its importance, redefining stock boundaries remains challenging due to the inherent complexities in accurately delineating natural populations (Cadrin et al., 2023).

Genetic markers have become essential for examining population structure and delineating fish stocks, enabling the identification of management units and assessing connectivity (Hauser & Carvalho, 2008; Ovenden et al., 2015). Within this context, single nucleotide polymorphisms (SNPs) have gained prominence due to their high-number, genome-wide distribution, and suitability for high-throughput genotyping (Seeb et al., 2011). The advent of cost-efficient SNP discovery and genotyping methods, such as reduced representation DNA sequencing (RAD-seq) (Catchen et al., 2017) or low coverage whole genome sequencing (lcWGS) (Therkildsen & Palumbi, 2017), has enhanced the capacity to detect subtle population differentiation, a feature that is particularly valuable for marine fishes, characterized by high dispersal potential and exposure to limited physical barriers (Gagnaire et al., 2015). However, despite recent efforts employing genome-wide SNP markers to elucidate its population structure and connectivity, the European anchovy (*Engraulis encrasicolus*) continues to exemplify a species for which stock boundaries remain contentious and unresolved.

The European anchovy is a commercially important small pelagic fish distributed along the eastern Atlantic, from Norway to South Africa, as well as in the Mediterranean and Black Sea (Reid, 1966). Along the Atlantic European coastline, management of this species has historically been structured around two administratively defined stocks: the Bay of Biscay stock (ICES Subarea 8) and the Atlantic Iberian Waters stock (ICES Division 9.a), with regions further north remaining unassessed (ICES, 2024). Notably, there is evidence indicating that the Gulf of Cadiz, the southernmost area of the Atlantic Iberian Waters stock (9.a. South), hosts a stable population that is largely independent from more northerly areas (9.a. West) (Ruiz et al., 2009). This independence is reflected in divergent biomass trends observed between both areas within 9.a, which prompted ICES to produce separate management advice for each of them from 2018. Nevertheless, a single TAC for Division 9a, comprising the aggregate advice for both components, was set by managers. Acknowledging that this approach could potentially lead to overexploitation of the component with limited fishing opportunities and, drawing upon a comprehensive synthesis of existing data (Garrido et al., 2024), ICES has, from 2024 onwards, issued separate advice for for 9a.Sout and 9a.West..

Over the last decades, numerous studies have investigated the population structure of European anchovy at different geographical scales using a variety of genetic markers, including allozymic variants (Sanz et al., 2008), mitochondrial DNA (Borrell et al., 2012; Magoulas et al., 2006; Silva, Horne, et al., 2014; Viñas et al., 2014), microsatellites (Borrell et al., 2012; Silva, Horne, et al., 2014; Zarraonaindia et al., 2009) and, more recently, Single Nucleotide Polymorphisms (Catanese et al., 2017; Huret et al., 2020; Montes et al., 2016; Petitgas et al., 2012; van der Kooij et al., 2024; Zarraonaindia et al., 2012). Despite these efforts, results regarding connectivity have often been inconclusive or contradictory, which was due to some studies being based on insufficient sample coverage and others not accounting for the complex evolutionary history of European anchovy, which encompasses the presence of genetically distinct co-occurring ecotypes: an “offshore” or “marine” ecotype distributed over the continental shelf, and an “inshore”, “estuarine” or “coastal” ecotype restricted to estuaries or their plumes (hereafter referred to as marine and coastal, respectively) (Bonhomme et al., 2022; Bouchenak-Khelladi et al., 2008; Catanese et al., 2017; Huret et al., 2020; Karahan et al., 2014; Le Moan et al., 2016; Montes et al., 2016; Oueslati et al., 2014). Notably, genetic differentiation between these two ecotypes exceeds than that observed among geographically distant locations within the same ecotype (Catanese et al., 2017; Le Moan et al., 2016). Further complicating this scenario, both ecotypes are known to co-occur and hybridize, although the extent of admixture remains undetermined (Le Moan et al., 2016; Montes et al., 2016).

Regarding the connectivity between the individuals from the Bay of Biscay stock and the individuals from the Atlantic Iberian Waters stock, studies based on a small number of markers and not considering ecotypes result in inconclusive outcomes, suggesting presence of multiple groups (Silva, Horne, et al., 2014; Zarraonaindia et al., 2012) and genetic heterogeneity between different locations within the Iberian Waters stock (Sanz et al., 2008; Borrell et al., 2012; Zarraonaindia et al., 2009; Zarraonaindia et al., 2012). There are also some doubts regarding the connectivity between the stocks assessed by ICES and populations from northernmost and southernmost locations. Several studies suggest that the Bay of Biscay is genetically distinct from northern areas (North Sea, Irish Sea and English Channel) (Huret et al., 2020; Montes et al., 2016; Petitgas et al., 2012; van der Kooij et al., 2024; Zarraonaindia et al., 2012), whereas few analyses have tackled the connectivity with north African Atlantic locations (Catanese et al., 2017; Montes et al., 2016; Silva, Horne, et al., 2014; Zarraonaindia et al., 2012).

More recently, studies based on whole genome sequencing data have provided new insights into anchovy population structure and its complex evolutionary history. Combining whole genome and restriction site associated DNA sequencing data of 39 and 385 samples respectively, Meyer et al. (2025) revealed a broad-scale structure in the Northeast Atlantic shaped by three major genetic ancestries, including two ecotypes —marine, coastal, and a southern lineage. Using whole genome sequencing data of 40 individuals Pujolar et al. (2025) found three genetically differentiated regional clusters, the first corresponding to marine anchovies from the Gulf of Cadiz, the second with individuals from the Kattegat Bay, and the third with individuals from Ireland and western Portugal. Both studies coincide in finding most genetic diversity and adaptive potential concentrated within putative structural variants, highlighting the strong adaptive potential of the species. Despite these advances, the delineation of biologically meaningful stock boundaries in European anchovy is still not fully resolved. Uncertainty persists in the boundary between the two existing ICES stocks, and in understanding their connectivity with northern and southern peripheral regions, an aspect that is crucial for clarifying their potential role in the hypothesized northward expansion of the species. To address this gap, this study has compiled a dataset that includes thousands of genetic markers from individuals covering the two European stocks in the Atlantic as well as the regions to the north and south (from the English Channel, in the north, to Canary Islands, in the south, and including the Mediterranean Sea), as well as individuals from the two ecotypes. We used this comprehensive dataset to (1) clarify the relationships between marine and coastal ecotypes, to (2) assess whether management units align with biological groupings, and (3) investigate the connectivity between populations of the current ICES stocks and from neighboring northern and southern populations.

## 2. MATERIALS AND METHODS

A summarized schematic view of the samples included, and methods used in this study is shown in Figure S1.

### Sample collection, DNA extraction, and RAD-seq library preparation and sequencing

A total of 382 adult anchovies were sampled during different scientific surveys conducted along the northeast Atlantic coast, from the English Channel (north) to the Canary Islands (south), including the Mediterranean Sea (Table S1, Figure 1A). From each anchovy, a ∼1 cm^3^ piece of muscle tissue was excised and immediately stored in RNA-later or 96% molecular grade ethanol at −20°C until DNA extraction. Genomic DNA extraction was performed using the Wizard® Genomic DNA Purification Kit (Promega), starting from 20 mg of tissue and following the manufacturer’s instructions. Extracted DNA was eluted in sterile Milli-Q water and its concentration was determined with Quant-iT dsDNA HS assay kit using a Qubit® 2.0 Fluorometer (Life Technologies). DNA integrity was assessed by electrophoresis, migrating about 100 ng of GelRed™-stained DNA on a 1.0% (w/v) agarose gel.

**Figure 1.**
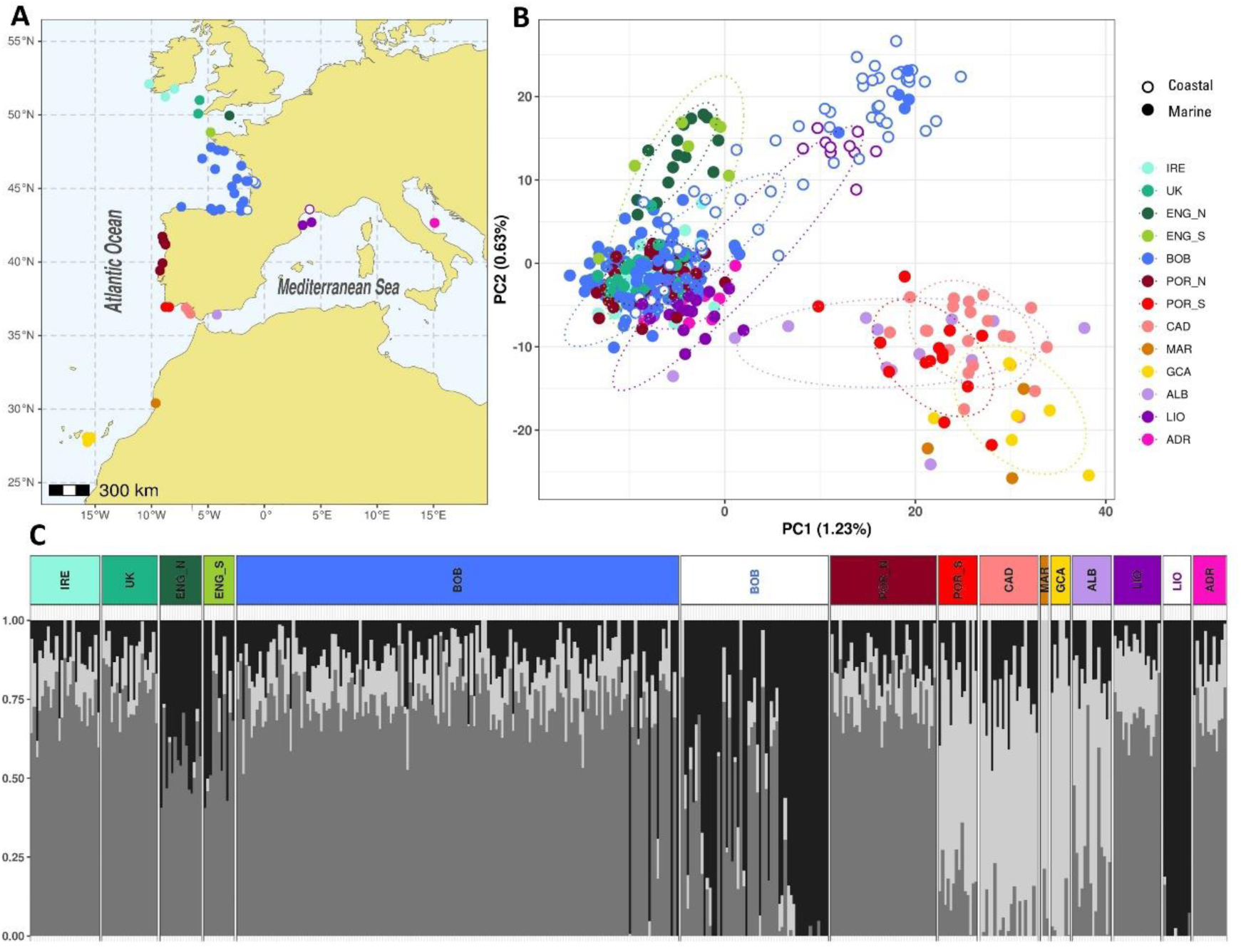
Population structure of the European anchovy based on neutral markers. **(A)** Map showing the catch location of the individuals included in this study. Each colour represents a sampling area; filled and open circles correspond to marine or coastal locations respectively. **(B)** Principal component analysis (PCA) performed using all individuals (n=419, 6498 neutral SNPs). Each point represents one sample; colours and shapes as in A (IRE, Ireland; UK, United Kingdom; ENG_N, English Channel North; ENG_S, English Channel South; BOB, Bay of Biscay; POR_N, Portugal North; POR_S, Portugal South; CAD, Gulf of Cadiz; MAR, Morocco; GCA, Canary Islands; ALB, Alboran Sea; LIO, Gulf of Lion; ADR, Adriatic Sea). Ovals represent 95% inertia ellipses. **(C)** ADMIXTURE clustering approach based on all individuals, where each bar represents an individual and shade of grey its inferred membership to each of the three potential ancestral populations (K = 3). Labels coloured as in A.

RAD-seq libraries were prepared following the method described by Etter et al. (2011) with custom-modified adapters for NOVASEQ UDIs compatibility, based on the forward and reverse overhang sequences provided in the 16S library preparation protocol developed by Illumina. Briefly, 500 ng of starting DNA was digested with the *SbfI* restriction enzyme (Thermo Fisher) and ligated to modified Illumina P1 adapters containing 5-6 bp unique barcodes. Equimolar pools of DNA from 32 individuals were sheared using the Covaris® M220 Focused ultrasonicator ™ Instrument (Life Technologies) and size selected to 300–500 bp by AMPure XP Beads (Beckman) (Bronner et al., 2014). After Illumina P2 adaptor ligation, each library was amplified using 12 PCR cycles using Nextera XT primers with Dual Unique combinations. Each pool was paired-end sequenced (150 bp) on an Illumina NOVASEQ (Illumina Inc.).

### RAD-tag assembly and SNP calling

The above newly generated RAD-sequencing data were combined with compatible data from 128 anchovies from a previously published study (Le Moan et al., 2016) available in the Short Read Archive from GenBank (Bioproject PRJNA311981). The combined raw reads were analysed using Stacks version 2.65 (Catchen et al., 2013). De-multiplexing, exclusion of reads with an average Phred score lower than 28 or containing adaptor sequence, as well as truncating all reads to 95 nucleotides to remove low-quality bases at the sequence end was performed using the ‘process_radtags’ module; PCR duplicates were removed with the ‘clone_filter’ module. Reads were *de novo* assembled into putative loci using the ‘ustacks’ module with a minimum depth coverage (m) of 5 reads to form a stack, a maximum of 2 nucleotide mismatches (M) to form a putative locus, and a maximum of 4 mismatches to incorporate secondary reads. Only samples with more than 25000 putative loci were kept. A catalog of loci was generated using the ‘cstacks’ module, allowing a maximum 3 mismatches between sample loci (n). Matches of individual loci to the catalog were searched using the ‘sstacks’ module, and data were stored by locus and SNPs were called using the ‘tsv2bam’ and ‘gstacks’ modules respectively.

### SNP filtering and genotype table building

From the generated catalog, a genotype table including all individuals (n=502) was generated as follows. SNPs present in loci found in at least 75% of the individuals were selected and exported to PLINK and Variant Call Format (VCF) using the ‘populations’ module. Using PLINK version v1.90b (Purcell et al., 2007) and considering only SNPs called from forward reads, increasing threshold values for minimum genotyping rate for individuals and SNPs were applied to obtain a final genotype table with a minimum genotyping rate of 0.90 and 0.95 per individual and SNP respectively. To exclude from the analyses rare variants, non-informative for population structure analysis, or those potentially derived from sequencing or assembly errors, SNPs with minor allele frequency (MAF) lower than 0.05 were removed. Analyses of genetic relatedness of each possible pair of individuals was estimated using the relatedness function implemented in VCFtools (Danecek et al., 2011) and no related pairs were found. The dataset was pruned by selecting only the first SNP per tag and excluding SNPs in strong Linkage Disequilibrium (LD) using PLINK software (r^2^ threshold 0.2). The resulting overall dataset genotype table was exported to Structure and Genepop formats using PDGSpider 2.1.1.0 (Lischer & Excoffier, 2012). Outlier loci were identified using the multivariate analysis method implemented in the pcadapt R package (Luu et al., 2016), which does not require a prior grouping of the samples. A screeplot representing the percentage of variance explained by each PC was used to choose the number of principal components (K) following author recommendations, and SNPs with p-values (adjusted following Benjamini and Hochberg (1995)) below 0.05 were classified as outliers.

### Population structure analyses

The following analyses were performed on neutral and outlier markers separately. Principal component analysis (PCA) was performed using the adegenet R package (Jombart & Ahmed, 2011) to illustrate the main axes of genetic variation among individuals, with no previous population assignment of samples into groups. The number and nature of distinct genetic clusters were investigated using the model-based clustering method implemented in ADMIXTURE (Alexander et al., 2009) assuming from 2 to 7 ancestral populations (K) and setting 1000 bootstrap runs. A first ADMIXTURE run was launched for each value of K to check the number of steps necessary to reach the default 0.001 likelihood value during the first run. This information was used to set the ‘-c’ parameter (steps to be fulfilled in each bootstrapped run) that would assure convergence for each analysis (from 20 to 100 steps) for the bootstrapped runs. The value of K (ranging from 2 to 10) with the lowest associated error value was identified using ADMIXTURE’s cross-validation procedure.

## 3. RESULTS

### Three main genetically distinct lineages of European anchovy

Population structure analyses based on neutral markers revealed three main genetically distinct groups: a southern marine lineage, ranging from the Canary Islands to southern Portugal, a northern marine lineage, present mostly in northern Atlantic and Mediterranean marine locations and a coastal lineage found mostly in Atlantic and Mediterranean coastal or estuarine locations. This pattern was supported by both the principal component and the ADMIXTURE analyses (Figure 1B,C). The geographic distribution of the southern marine lineage included the Canary Islands, Marocco, the south of Portugal and the Gulf of Cadiz but excluded Northern Portugal. Within this lineage, individuals from Marocco and the Canary Islands appeared to be partially differentiated from individuals from Cadiz and the south of Portugal. Additionally, the Alboran Sea was noted as a transition zone, harbouring individuals clustering within the southern lineage, individuals clustering within the marine lineage and individuals that appeared to be hybrids. The northern marine lineage consisted of marine individuals present in the Atlantic Ocean and Mediterranean Sea, which were genetically more closely related among themselves than to those of the southern lineage. Within this group, the English Channel was genetically distinct from the rest of the northeast Atlantic locations, and the Mediterranean and Atlantic locations could only be clearly distinguished by plotting the first and third principal components (Figure S2). The coastal lineage consisted mostly of individuals sampled from coastal or estuarine locations. Genetic analyses revealed that, despite being geographically isolated, samples from estuaries in the Atlantic, and a lagoon in the Mediterranean were genetically more similar to each other than to samples from marine locations nearby (Figure 1B,C) and that, within the locations for which both ecotypes were available, marine samples were closer to each other than coastal samples (Figure S3). Despite the coastal and marine lineages being genetically differentiated, they coexist, yet at different space-time proportions. The analyses showed the presence of individuals from both genetic groups within the estuary and near the estuary, as well as of genetically intermediate individuals, indicating that hybridization occurs between the two groups. Given that about one third of the SNPs of the dataset were either in Linkage disequilibrium (1995 SNPs) or identified as outliers (716 SNPs), we also performed analyses based on them. Overall, results show the same main connectivity pattern as when using neutral markers (Figure S4), suggesting that the results observed are due to neutral processes rather to adaptation.

### Mismatch between stock and genetic boundaries in the ICES assessed marine lineage

To better understand the connectivity of European anchovy populations in a stock boundary context, we performed genetic analysis excluding the coastal lineage (Figure 2). Results showed that the pattern of genome-wide marker-based population structure defined for the northeastern Atlantic anchovy did not align with ICES defined stocks. The Bay of Biscay stock (ICES Subarea 8) was genetically connected to the north with the Celtic Seas, an area outside ICES stock boundary, and to the south was connected to the northwestern Iberia, which corresponds to the northernmost area of the Atlantic Iberian anchovy stock. Individuals from the English Channel (except for some in the southernmost section of the Channel) were genetically distinct of those from the Bay of Biscay stock and Celtic Seas (also seen in Figure 1B,C). The Atlantic Iberian Waters stock (ICES Division 9a) was shown to contain two genetically distinct groups which in turn are connected to other areas: (1) the northernmost area is connected to the Bay of Biscay stock and Celtic Seas (outside ICES stock boundaries) and (2) the southernmost area is connected to the African coast (outside ICES stock boundaries), and to a lesser extent, to the Mediterranean Sea by the Alboran sea.

**Figure 2.**
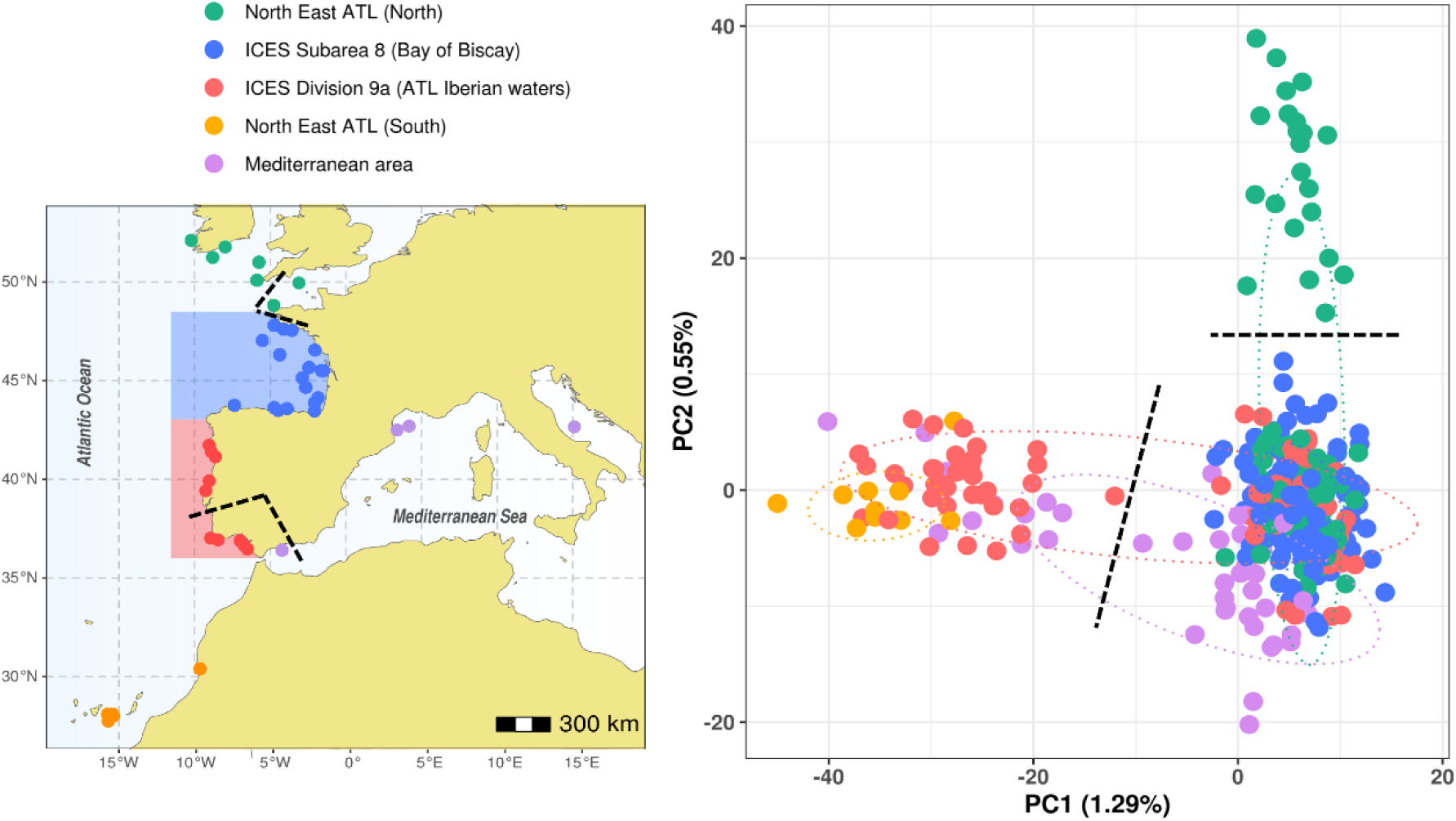
Distribution of genetically differentiated anchovy clusters among stocks. **(A)** Map showing the catch location of the anchovy individuals, where colours indicate Bay of Biscay stock – blue, Atlantic Iberian waters stock – red, English Chanel – green, north African coast and Canary Islands – orange, Mediterranean – purple. **(B)** PCA performed without the coastal lineage. For clarity, lines showing where the PCA breaks align with the map have been included.

## 4. DISCUSSION

This study addresses previously unresolved issues in European anchovy population genetic structure and provides new insights into the connectivity patterns of this species in European and North African waters. We identify three main genetic lineages: a northern marine lineage, a southern marine lineage, and a coastal lineage, suggesting that the evolutionary history of the European anchovy is more complex than previously thought. This explains conflicting or unconclusive results derived from studies that tried to explain patterns in the northern marine lineage, without or only partially accounting for the presence of two additional linages (e.g. Borrell et al. (2012); Catanese et al. (2017); Huret et al. (2020); Le Moan et al. (2016); Magoulas et al. (2006); Montes et al. (2016); Petitgas et al. (2012); Sanz et al. (2008); Silva, Horne, et al. (2014); van der Kooij et al. (2024); Viñas et al. (2014); Zarraonaindia et al. (2012); Zarraonaindia et al. (2009)). Building on recent genome-wide studies (Meyer et al., 2025; Pujolar et al., 2025), which revealed broad-scale genetic structure and the role of structural variants in anchovy lineage divergence, our study provides a finer-scale resolution by incorporating dense geographic sampling, inclusion of both marine and coastal ecotypes, and explicit evaluation of the alignment between genetic and management units. This approach allows us to confirm and expand upon previous findings while offering direct implications for fisheries management. In doing so, our study contributes to evolutionary theories of anchovy niche colonization, supports the reassessment of current stock boundaries, and provides a framework for anticipating future distributional changes.

### Influence of southern marine and coastal lineages on European anchovy connectivity studies

Our results revealed a southern marine lineage (with the northern limit located in south Iberian Atlantic waters of the Gulf of Cadiz), clearly differentiated from samples from western Iberia (north of Lisbon) and from the Mediterranean Sea. Previous studies have reported a genetic break in the Atlantic, around mid-Portugal (Silva, Horne, et al., 2014), or further North in Galicia (Silva, Lima, et al., 2014; Zarraonaindia et al., 2012). However, while these studies may have unknowingly captured different aspects of these complex admixed ancestries, leading to different and conflicting interpretations, the connectivity barrier we found between the two marine lineages located between mid-Portugal and the Gulf of Cadiz is consistent with recent results based on genome-wide genomic markers (Meyer et al., 2025; Pujolar et al., 2025). Additionally, we noted the Alboran Sea as a transition zone, with anchovies either genetically similar to the southern marine lineage, to the Mediterranean (part of the northern marine lineage) or hybrids. This challenges the presence of a connectivity barrier in the Almeria-Oran front as suggested by previous studies (Magoulas et al., 2006; Zarraonaindia et al., 2012). More recently, Meyer et al. (2025) described the Alboran Sea as a tripartite contact zone involving the marine, coastal, and southern lineages, with evidence of ancestry mixing. Our data support this interpretation, showing that individuals from multiple lineages coexist in the Alboran Sea, and suggesting that this region is a hotspot for admixture and the potential origin of novel genetic combinations.

In this context, our results also suggest that the coastal ecotype may have originated from the coastal linage. Presence of two ecotypes in European anchovy was previously described using SNP markers in North Western Atlantic locations, including the Gironde (Montes et al., 2016), Abra (Montes et al., 2016), Adour (Le Moan et al., 2016), Loire (Huret 2020) and Ijsselmer (Montes et al., 2016), and in the Mediterranean sea (Le Moan et al. (2016), Catanese et al. (2017), and Bonhomme et al. (2022)). All these studies show that coastal anchovies are genetically more similar with one another than to the neighbouring, and sometimes spatially overlapping, marine anchovies. Curiously, among the samples collected within this project, only one pure oceanic sample was detected within the Gironde estuary, and only four pure coastal anchovies (from 2023) were found outside the estuary, while data published by Le Moan et al. (2016) suggested much higher rates of habitat exchange and hybridization in the Adour estuary. This suggests that the distribution of both ecotypes could be dynamic and subjected to temporal and/or geographical changes and calls for a continuous monitoring of their spatiotemporal distribution.

Within the Bay of Biscay, the absence of genetic differences observed among locations contrasted with the heterogeneity reported previously in studies using allozymes (Sanz et al., 2008), microsatellites (Borrell et al., 2012; Zarraonaindia et al., 2009), mitochondrial DNA (Borrell et al., 2012) and SNP markers (Zarraonaindia et al., 2012). The heterogeneity found in these previous studies could be explained by the fact that they did not consider the presence of different ecotypes in the area.

### A shared northern marine lineage between the Atlantic and Mediterranean populations

Our results show that, although clearly differentiated, individuals from the northern Atlantic marine locations are genetically more closely related to the Mediterranean marine locations than to those of the southern or coastal lineages. These results support the hypothesis of a common ancestry of the northern marine lineage, as evidenced by previous studies suggesting a common ancestor for Bay of Biscay and western Mediterranean anchovy populations (Catanese et al., 2017; Magoulas et al., 2006; Montes et al., 2016; Zarraonaindia et al., 2012; Zarraonaindia et al., 2009).

While this supports a broad genetic continuity between the northern Atlantic and Mediterranean marine populations, our results also reveal finer-scale structure within the northern Atlantic, (particularly in the English Channel and Celtic Sea). Our population structure analyses show that the English Channel is genetically distinct from the rest of the northeast Atlantic locations. This consolidates the genetic differentiation found in previous studies (Huret et al., 2020; Montes et al., 2016; Petitgas et al., 2012; van der Kooij et al., 2024; Zarraonaindia et al., 2012) according to which the Bay of Biscay individuals are genetically distinct from the northernmost populations (the North Sea, Irish Sea and the English Channel).

Interestingly, Huret et al. (2020) showed that, in autumn, the spatial ranges of part of Bay of Biscay and English Channel anchovy populations slightly overlap in the west of Britanny, Celtic Sea region. In our study, anchovy individuals from the English Channel were collected in autumn, which could explain why some individuals collected in the southernmost section of the Channel belong to the Bay of Biscay stock individuals while others grouped with northernmost populations. Unlike previous studies that included samples from the North Sea (Huret et al., 2020; Montes et al., 2016; Petitgas et al., 2012; Silva, Horne, et al., 2014; van der Kooij et al., 2024; Zarraonaindia et al., 2012) and the Irish Sea (Huret et al., 2020; Montes et al., 2016; van der Kooij et al., 2024), our study does not cover those areas. These previous studies consistently reported that anchovy individuals from the North Sea and Irish Sea belong to a northern genetic cluster that is clearly distinct from the Bay of Biscay population. However, we included samples from the Celtic Sea (UK and Ireland), a region that had not been genetically characterized in earlier work. Interestingly, individuals from these locations consistently grouped with the Bay of Biscay lineage, suggesting genetic continuity between the Bay of Biscay and the Celtic Sea. This finding expands the known northern range of the Bay of Biscay genetic group and highlights the importance of including the Celtic Sea region in future assessments of population structure.

### Implications for management

Our results reveal that the genome-wide marker-based population structure of northeastern Atlantic anchovy does not fully match the current ICES stocks boundaries. This mismatch has important implications for an effective management and long-term conservation of the species.

First, the Bay of Biscay stock extends beyond the current administrative boundaries used for management. Our findings indicate connectivity between the Bay of Biscay stock (ICES Subarea 8) with the Celtic Sea, along with partial separation from the English Channel, which might act as a transition zone. This pattern is consistent with the findings of Huret et al. (2020), who also reported seasonal differences, with distribution overlap observed between more northern locations and the Bay of Biscay stock in autumn, and reduced overlap in spring and summer. The recognition of this extended boundary has been partially incorporated into the Bay of Biscay stock assessment, which allocates catches from statistical rectangles 25E4 and 25E5 within subarea 7 that are adjacent to Subarea 8 to the Bay of Biscay stock (ICES, 2025a, 2025b). The variability detected in the English Channel, where some samples cluster genetically with the Bay of Biscay stock while others are distinct, underscores the need for further analyses, including consideration of seasonal dynamics, to guide the allocation of catches from this transitional zone. Such analyses would support the establishment of a flexible or shared management area that reflects the dynamic and seasonal distribution patterns of anchovy. Notably, van der Kooij et al. (2024), reported a northward expansion of the Bay of Biscay anchovy stock in 2019-2020, with post-larval individuals reaching the English Channel. This expansion was not observed in 2021, suggesting that such may be episodic and influenced by environmental and demographic factors. These changes affect stock management by potentially misaligning survey coverage and fishing effort.

Second, the Bay of Biscay stock exhibit genetic connectivity with the northern sector of the Atlantic Iberian Waters stock (ICES Division 9a), with a genetic discontinuity observed between Northern and Southern Portugal. These findings, which align with recent genome-wide analyses (Meyer et al., 2025; Pujolar et al., 2025), advocate for a redefinition of the European anchovy stock boundaries, suggesting a new demarcation south of Lisbon. This proposed revision is further substantiated by stable isotope analysis of the anchovy eye lenses (Sakamoto and Garrido, submitted) and by oceanographic modelling studies (e.g., Teles-Machado et al. (2024)), which identify larval dispersal from the Bay of Biscay to Portuguese waters but not beyond mid-Portugal, as well as lack of evidence of larvae from the Gulf of Cadiz dispersing northward. Collectively, these findings highlight the biological significance of the proposed boundary and provide strong support for the genetic isolation of the southern stock. Notably, preliminary findings from our genetic study were included in a comprehensive working document by Garrido et al. (2024), which was presented and evaluated during the ICES benchmark for anchovy stocks (ICES WKBANSP 2024). Following this benchmark, the anchovy stock in Division 9a was redefined, resulting in its separation into two distinct units: the southern component of Division 9a (encompassing the Gulf of Cadiz and the southern coast of Portugal) and the western component of Division 9a (covering the western Iberian waters).

Third, our results confirm that the two genetically distinct ecotypes, marine and coastal, coexist, within river plumes and adjacent coastal areas. However, we found a low number of coastal individuals in samples collected during offshore surveys. Consistent with Le Moan et al. (2016) and Meyer et al. (2025), we also identified hybrids between the two ecotypes, which highlights the need for more research into what maintains these differences and if they are due to environmental adaptation. While the short-term influence of the coastal ecotype on anchovy biomass estimates seems minor, it should still be considered, as its frequency in regions near river mouths might vary. Future assessments could benefit from monitoring ecotype admixture and migration rates and determining if correction factors are necessary in specific contexts. Ongoing genetic monitoring could further reveal whether coastal ecotypes serve as demographic sinks or sources for marine populations, which has implications for resilience and adaptive potential under environmental change.

### Conclusions and Future Directions

Our research provides strong evidence supporting a revision of the current International Council for the Exploration of the Sea (ICES) boundaries for European anchovy. The genetic data presented here show that existing stock definitions do not accurately represent actual biological populations and help clarify earlier uncertainties regarding anchovy population structure and connectivity. Additionally, the discovery of many outlier genetic markers highlight strong signals of local adaptation, which may maintain genetic differences despite notable gene flow. Analyses using both these adaptive markers and a set of neutral markers filtered for high linkage disequilibrium reveal consistent patterns of population structure, supplying robust proof of genuine demographic isolation among the three lineages identified in this study. These findings are crucial for sustainable harvest of the European anchovy, a species of significant ecological and economic value. Furthermore, the set of single nucleotide polymorphisms (SNPs) identified here provides a valuable base for future genetic monitoring tools, which could aid in catch assignment and enhance the management of anchovy fisheries by allowing more accurate identification of ecotype or stock origin.

## 5. ACKNOWLEDGEMENTS

We thank Guillermo Boyra, Ciaran O’Donnell, Fernando Ramos, Pablo Carrera, Antón Urbano, Ana Moreno, Erwan Duhamel, Jeroen Van der Kooij, Jose Pajuelo, Marta Coll, Federico Cali and Marie-Laure Acolas for providing the samples used in this study, and Iñaki Mendibil and Natalia Gutiérrez for excellent laboratory work. This study has been supported by Department of Food, Rural Development, Agriculture and Fisheries of the Basque Government through the GENGES project; by the EMFF (European Maritime and Fisheries Fund) Data Collection Framework through the MPDH project; and by the General Fisheries Secretariat of the Spanish Government, which provided access to the oceanographic vessels *Emma Bardán* and *Vizconde de Eza*.

## 7. SUPPLEMENTARY TABLE

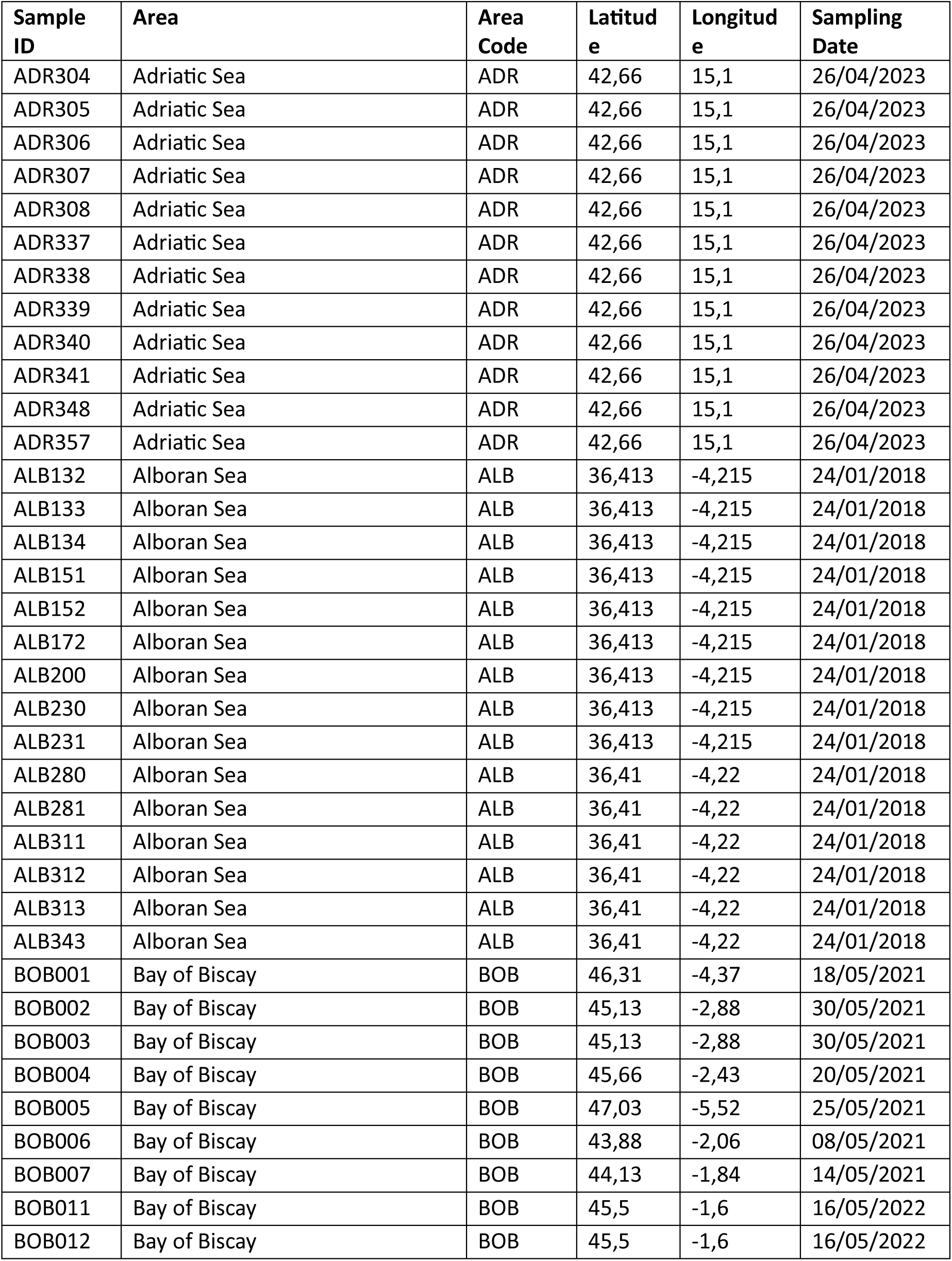
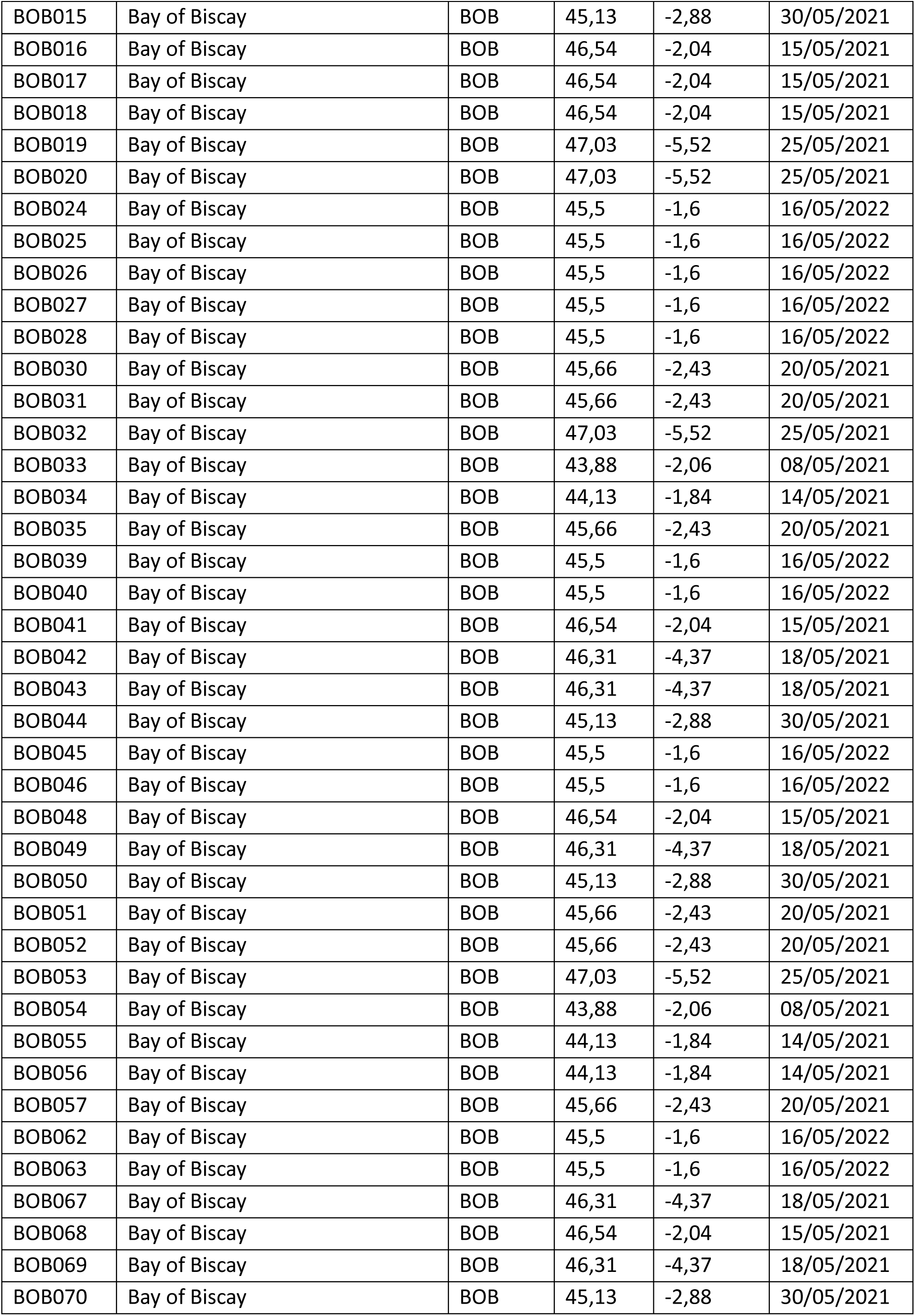
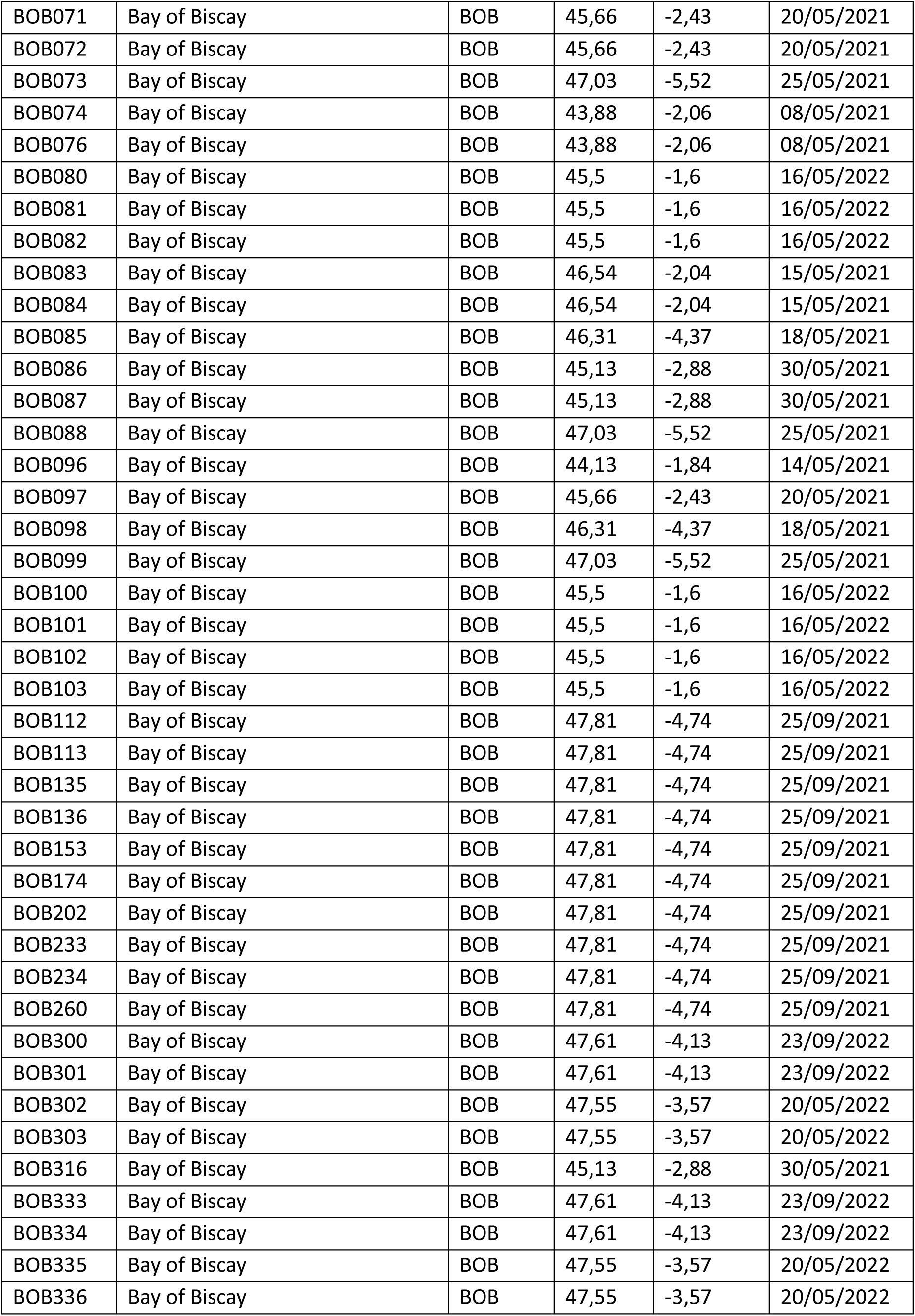
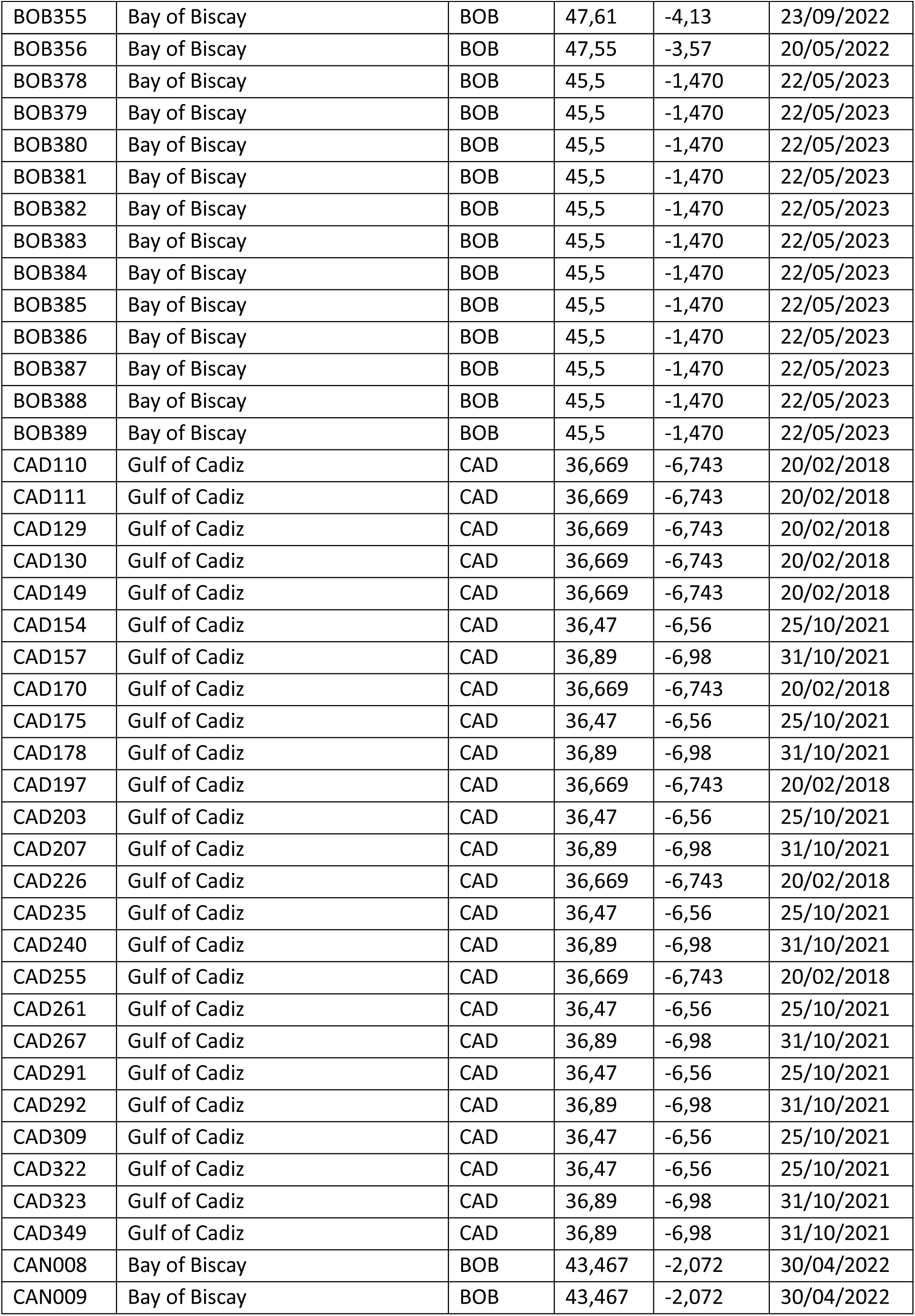
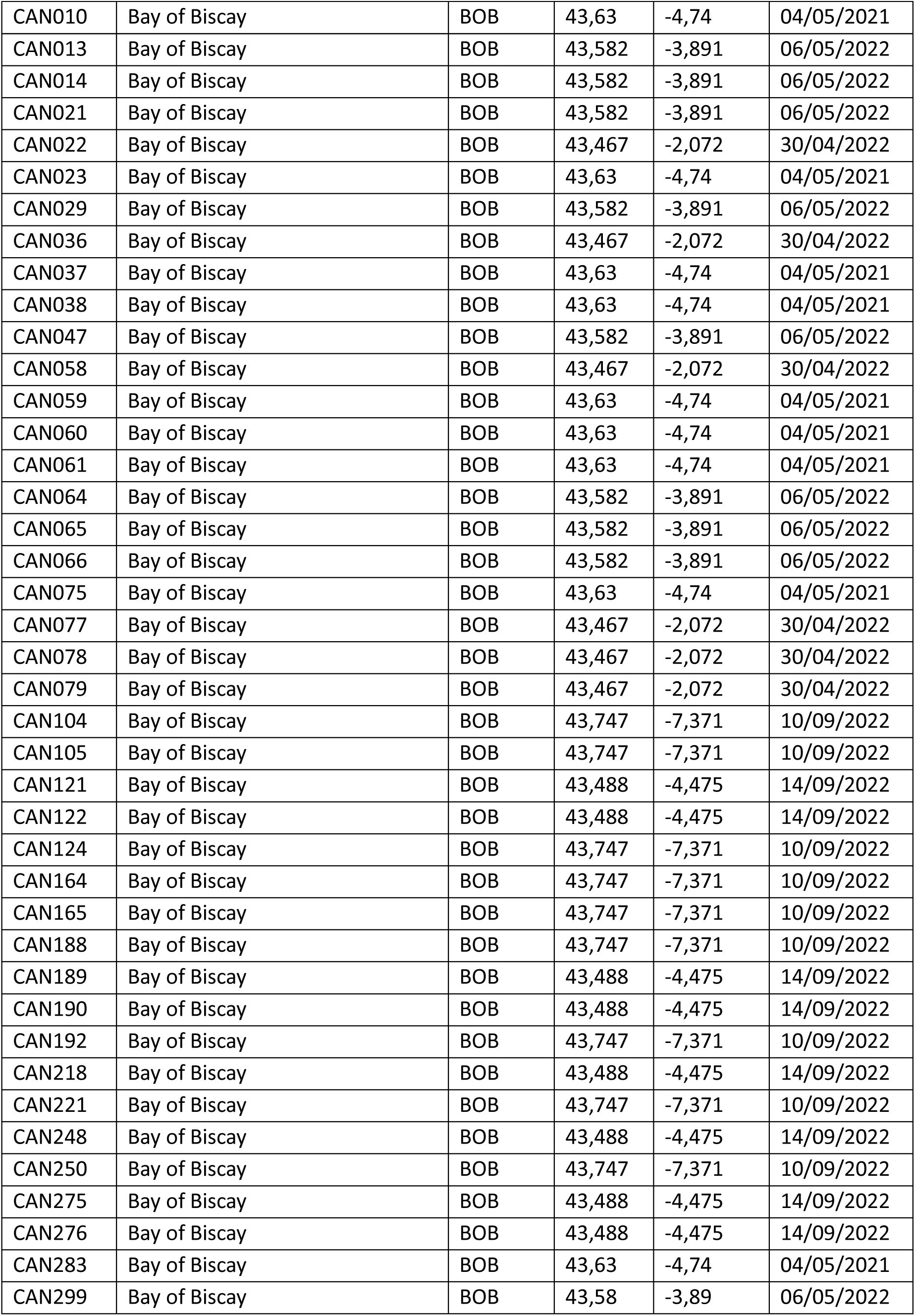
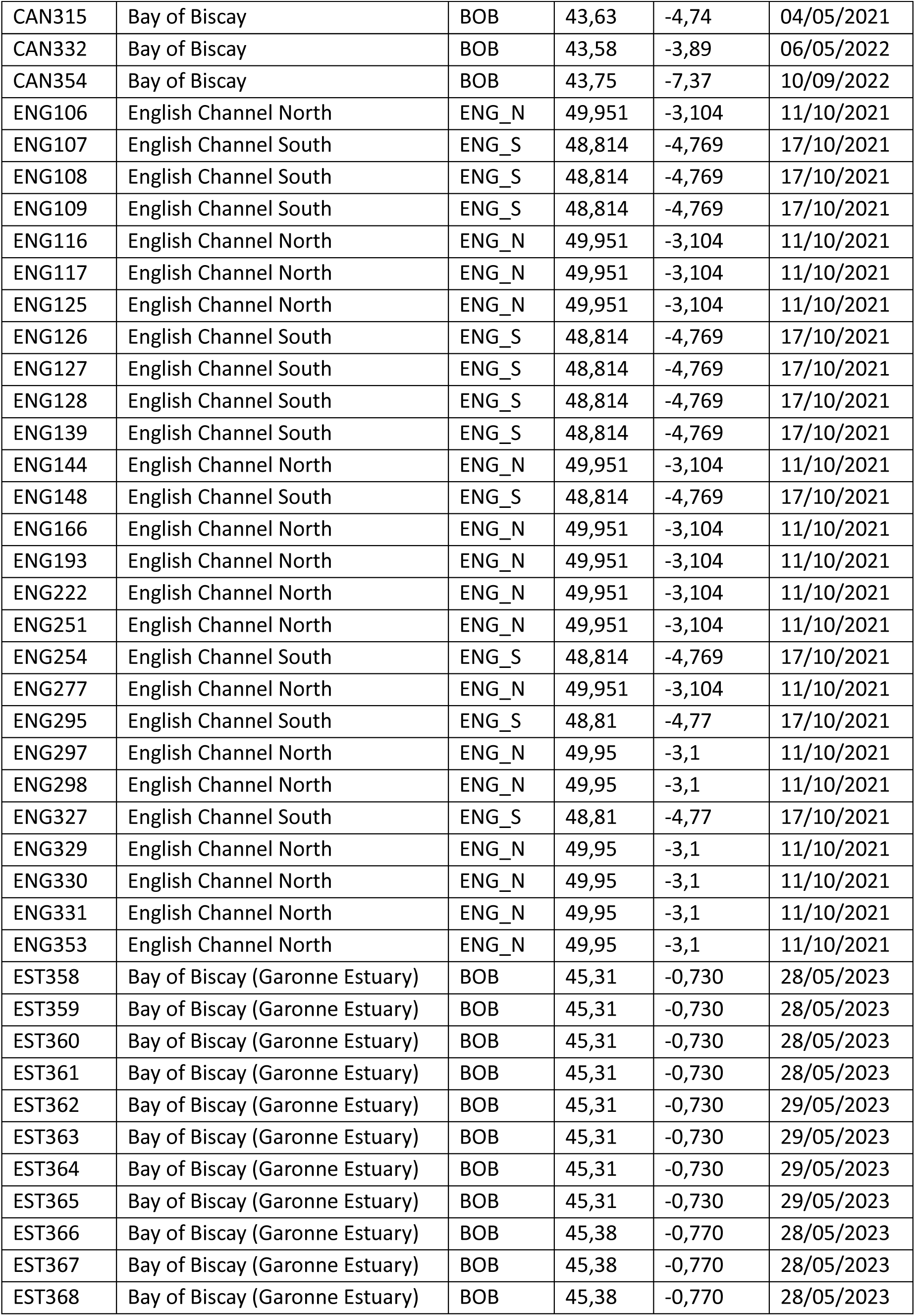
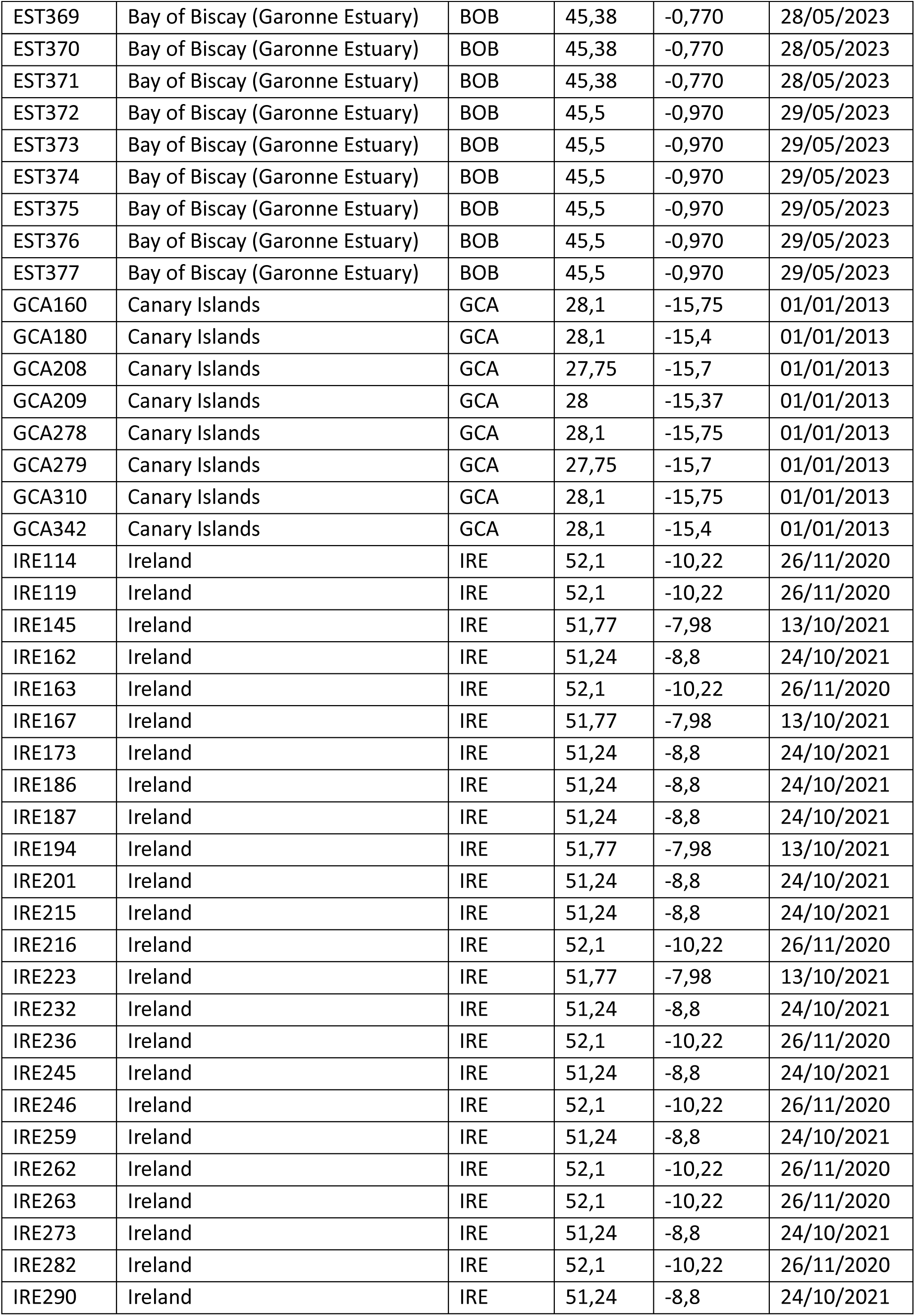
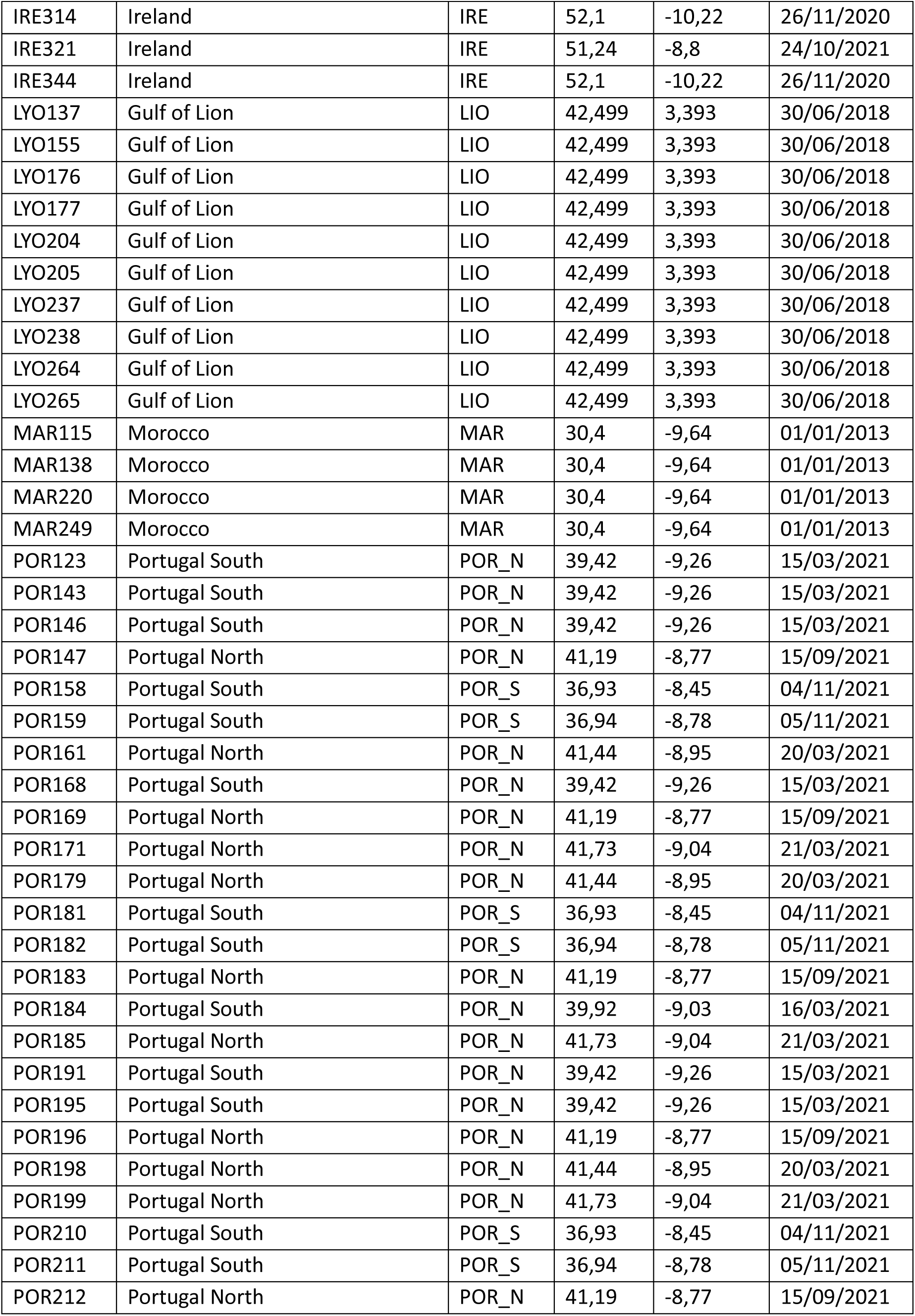
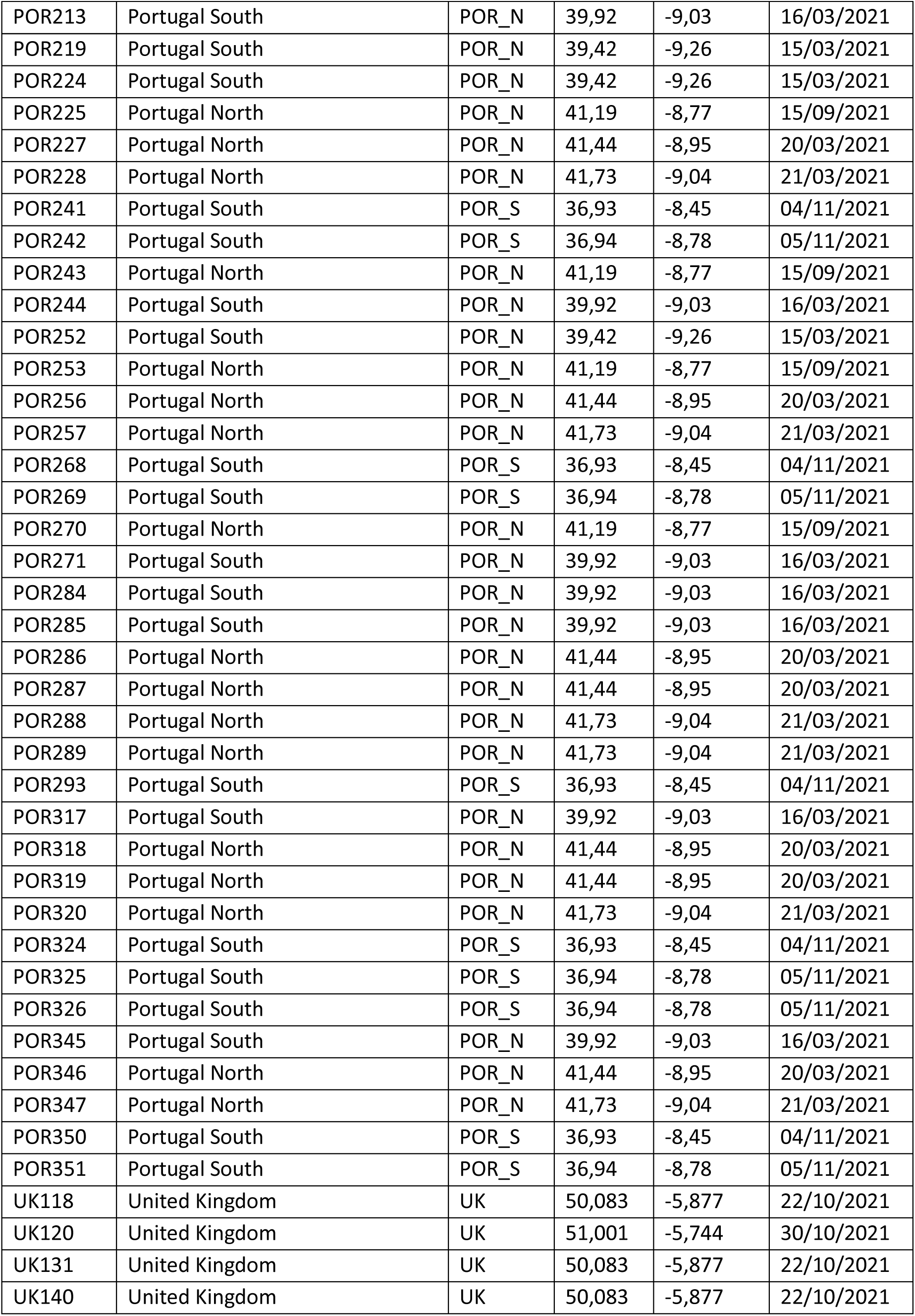
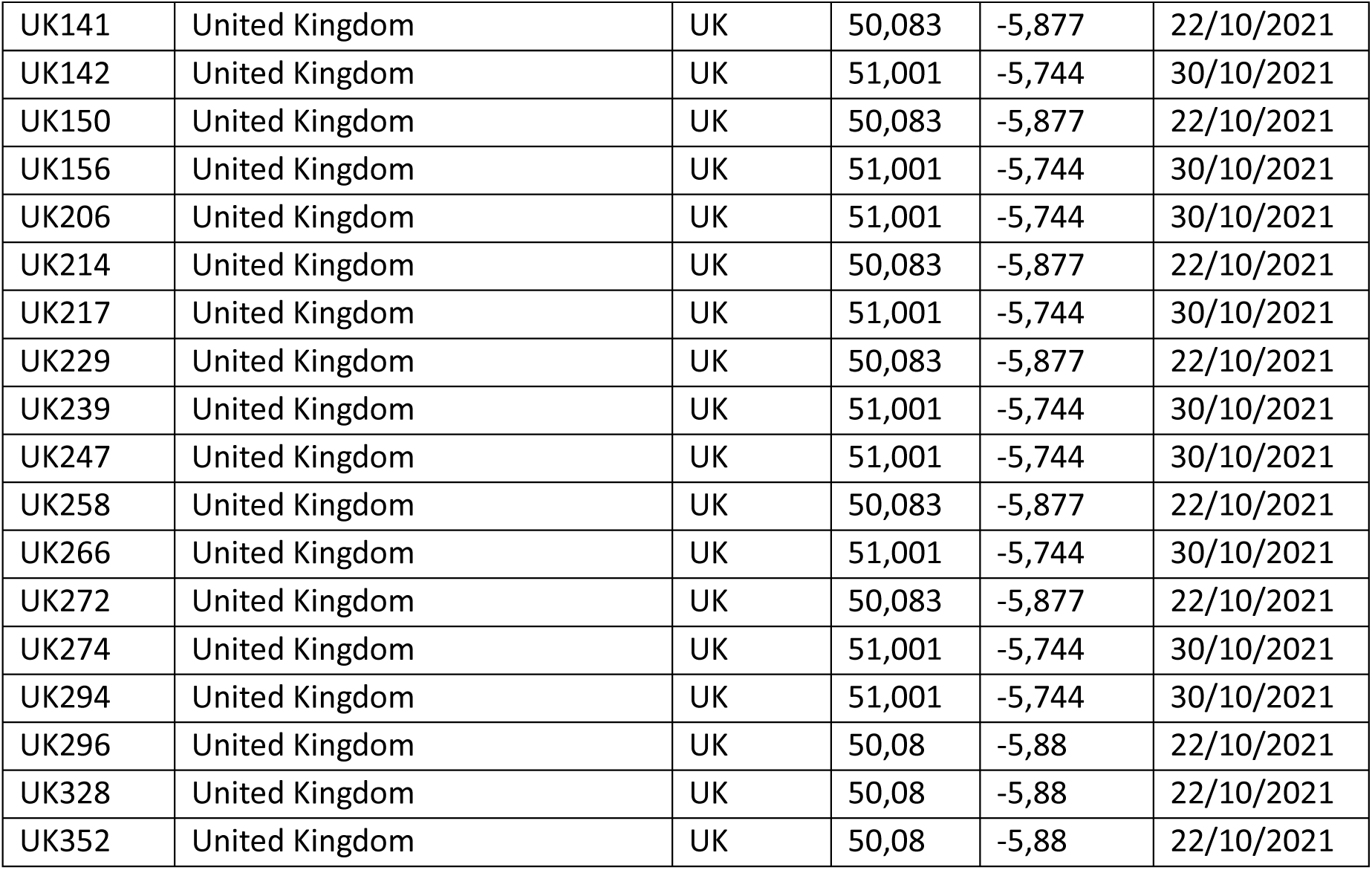

## 8. SUPPLEMENTARY FIGURES

**Figure S1.**
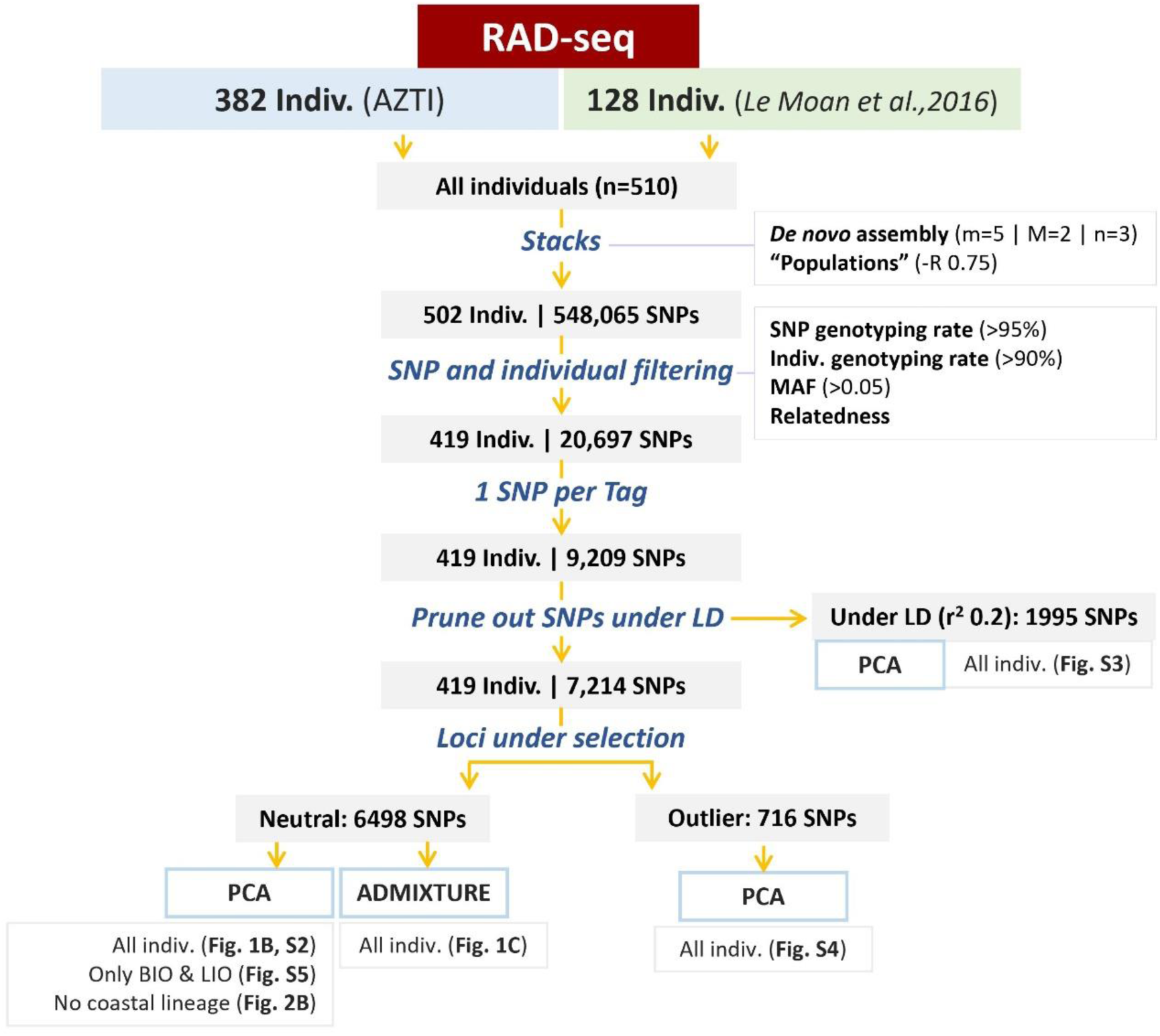
Detailed schematic view of the workflow done in this study.

**Figure S2.**
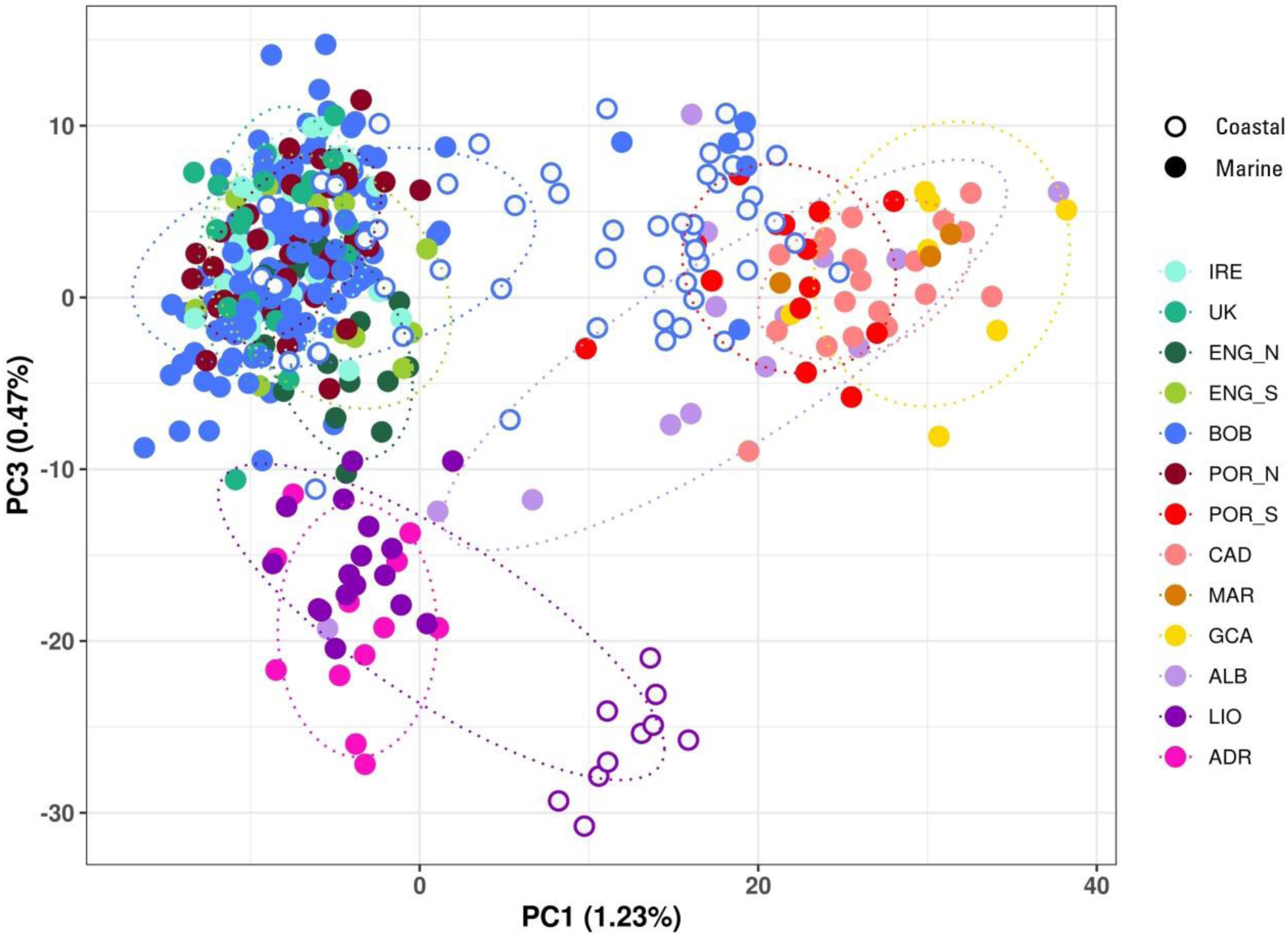
Principal component analysis (PCA) performed using PC1 and PC3 (all individuals, n=425, 6498 neutral SNPs). Each point represents one sample; colours denote the sampling area, and filled and open circles correspond to marine or coastal locations respectively (colours as in Figure 1). Ovals represent 95% inertia ellipses.

**Figure S3.**
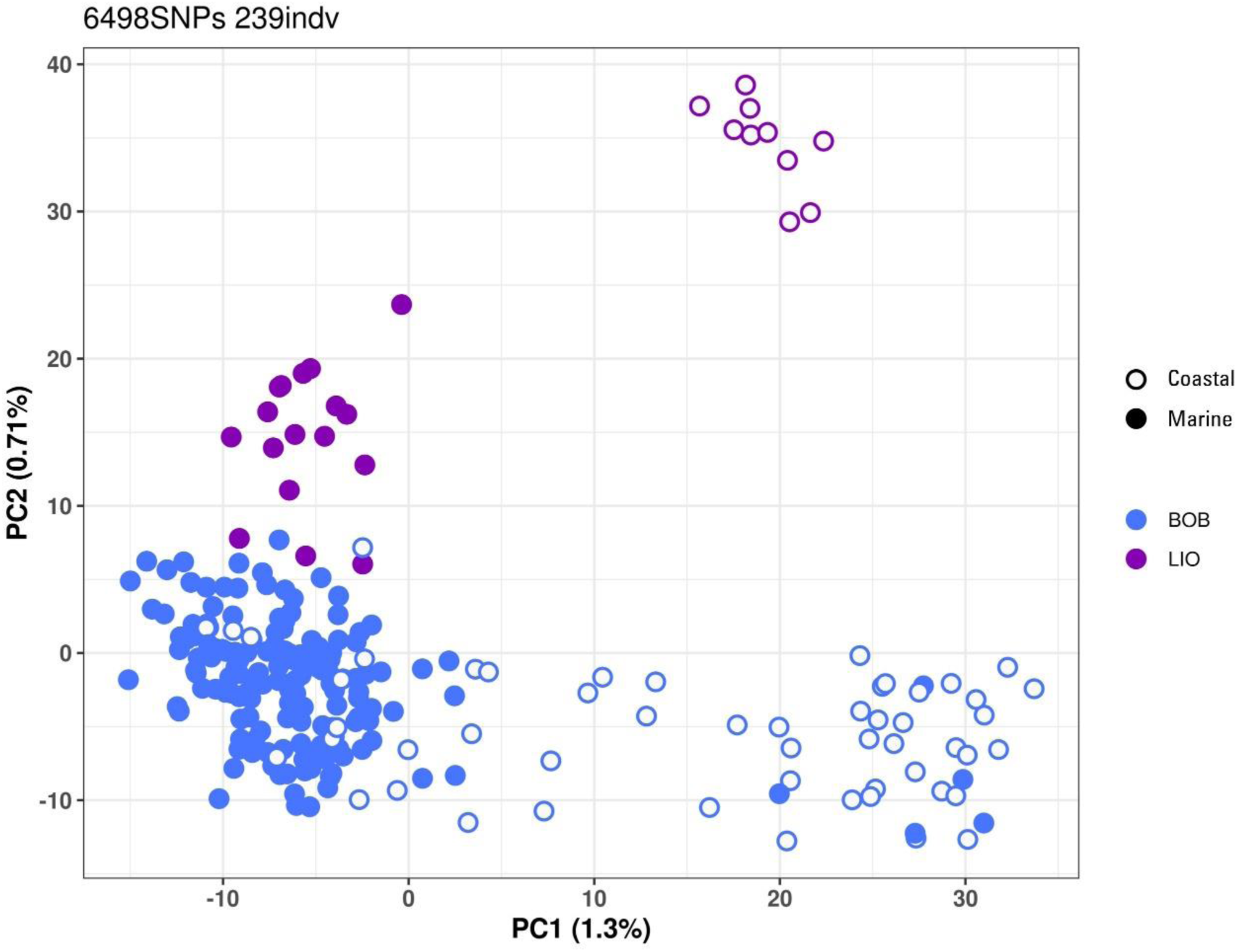
Principal component analysis (PCA) performed using marine (filled circle) and coastal (open circle) individuals collected around the Bay of Biscay (BIO) and the Gulf of Lion (LIO) areas (n=239, 6498 neutral SNPs). Each point represents one sample.

**Figure S4.**
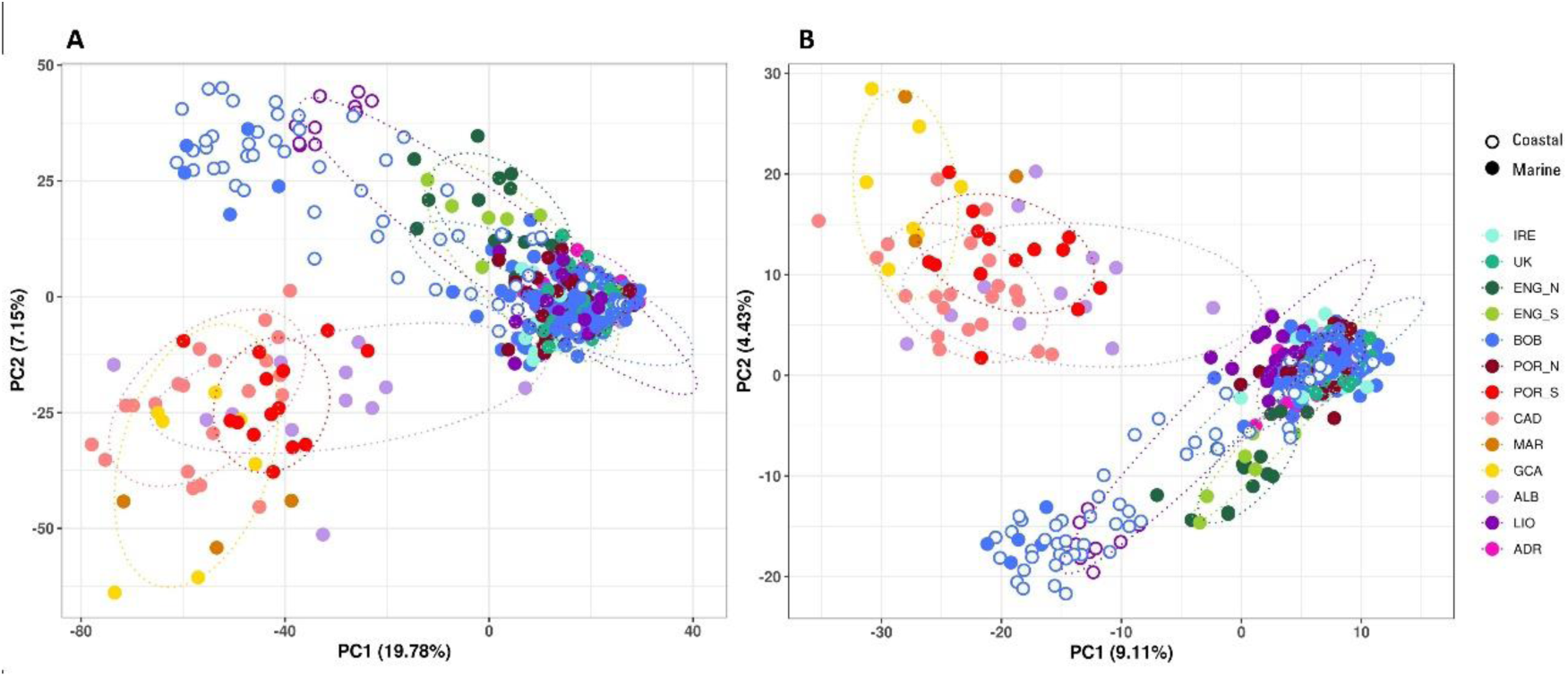
Principal component analysis (PCA) performed using all individuals (n=419). (A) Using 1995 SNPs under Linkage Disequilibrium. (B) Using 716 SNP identified as outliers by ‘pcadapt’. Each point represents one sample; colours denote the sampling area, and filled and open circles correspond to marine or coastal locations

## BIBLIOGRAPHY

Benjamini, Y., & Hochberg, Y. (1995). Controlling the False Discovery Rate: A Practical and Powerful Approach to Multiple Testing. J R Statist Soc, 57(1), 289–300. 10.2307/2346101

Bonhomme, F., Meyer, L., Arbiol, C., Banaru, D., Bahri-Sfar, L., Fadhlaoui-Zid, K., Strelkov, P., Arculeo, M., Soulier, L., Quignard, J. P., & Gagnaire, P. A. (2022). Systematics of European coastal anchovies (genus Engraulis Cuvier). J Fish Biol, 100(2), 594–600. 10.1111/jfb.14964

Borrell, Y. J., Piñera, J. A., Sánchez Prado, J. A., & Blanco, G. (2012). Mitochondrial DNA and microsatellite genetic differentiation in the European anchovy Engraulis encrasicolus L. ICES Journal of Marine Science, 69(8), 1357–1371. 10.1093/icesjms/fss129

Bouchenak-Khelladi, Y., Durand, J.-D., Magoulas, A., & Borsa, P. (2008). Geographic structure of European anchovy: A nuclear-DNA study. Journal of Sea Research, 59(4), 269–278. 10.1016/j.seares.2008.03.001

Bronner, I. F., Quail, M. A., Turner, D. J., & Swerdlow, H. (2014). Improved Protocols for Illumina Sequencing. Curr Protoc Hum Genet, 80, 1–42. 10.1002/0471142905.hg1802s80

Cadrin, S. X., Goethel, D. R., Berger, A., & Jardim, E. (2023). Best practices for defining spatial boundaries and spatial structure in stock assessment. Fisheries Research, 262, 106650. 10.1016/j.fishres.2023.106650

Catanese, G., Watteaux, R., Montes, I., Barra, M., Rumolo, P., Borme, D., Buongiorno Nardelli, B., Botte, V., Mazzocchi, M. G., Genovese, S., Di Capua, I., Iriondo, M., Estonba, A., Ruggeri, P., Tirelli, V., Caputo-Barucchi, V., Basilone, G., Bonanno, A., Iudicone, D., & Procaccini, G. (2017). Insights on the drivers of genetic divergence in the European anchovy. Scientific Reports, 7(1), 4180. 10.1038/s41598-017-03926-z

Catchen, J., Hohenlohe, P. A., Bassham, S., Amores, A., & Cresko, W. A. (2013). Stacks: an analysis tool set for population genomics. Molecular Ecology, 22(11), 3124–3140. 10.1111/mec.12354

Catchen, J. M., Hohenlohe, P. A., Bernatchez, L., Funk, W. C., Andrews, K. R., & Allendorf, F. W. (2017). Unbroken: RADseq remains a powerful tool for understanding the genetics of adaptation in natural populations. Molecular Ecology Resources, 17(3), 362–365. 10.1111/1755-0998.12669

Danecek, P., Auton, A., Abecasis, G., Albers, C. A., Banks, E., DePristo, M. A., Handsaker, R. E., Lunter, G., Marth, G. T., Sherry, S. T., McVean, G., Durbin, R., & Genomes Project Analysis, G. (2011). The variant call format and VCFtools. Bioinformatics, 27(15), 2156–2158. 10.1093/bioinformatics/btr330

Etter, P. D., Bassham, S., Hohenlohe, P. A., Johnson, E. A., & Cresko, W. A. (2011). SNP Discovery and Genotyping for Evolutionary Genetics Using RAD Sequencing. In V. Orgogozo & M. V. Rockman (Eds.), Molecular Methods for Evolutionary Genetics (Vol. 772, pp. 157–178). 10.1007/978-1-61779-228-1_9

Gagnaire, P.-A., Broquet, T., Aurelle, D., Viard, F., Souissi, A., Bonhomme, F., Arnaud-Haond, S., & Bierne, N. (2015). Using neutral, selected, and hitchhiker loci to assess connectivity of marine populations in the genomic era. Evolutionary Applications, 8(8), 769–786. 10.1111/eva.12288

Garrido, S., Rodríguez-Ezpeleta, N., Díaz, N., Machado, A., Sakamoto, T., Ramos, F., Rincón, M., Moreno, A., Jiménez, M. P., Santos, M., Carrera, P., Rodríguez-Climent, S., Feijó, D., Ibabarriaga, L., Citores, L., Boyra, G., & Duhamel, E. (2024). Population structure of the European Anchovy (Engraulis encrasicolus) in ICES Division 9a. Working document presented to the ICES Benchmark Workshop on Anchovy Species (WKBANSP), ICES Scientific Reports. 6: 96. 511 pp. 10.17895/ices.pub.27909783

Huret, M., Lebigre, C., Iriondo, M., Montes, I., & Estonba, A. (2020). Genetic population structure of anchovy (Engraulis encrasicolus) in North-western Europe and variability in the seasonal distribution of the stocks. Fisheries Research, 229, 105619. 10.1016/j.fishres.2020.105619

Hutchinson, W. F. (2008). The dangers of ignoring stock complexity in fishery management: the case of the North Sea cod. Biology Letters, 4(6), 693–695. 10.1098/rsbl.2008.0443

ICES. (2024). Working Group on Southern Horse Mackerel, Anchovy and Sardine (WGHANSA). ICES Scientific Reports, 6(46), 738. 10.17895/ices.pub26003356.v2

ICES. (2025a). Anchovy *Engraulis encrasicolus* in Subarea 8 (Bay of Biscay). ICES Stock Annexes, 47. 10.17895/ices.pub28053440.v1

ICES. (2025b). Benchmark Workshop for anchovy stocks (WKBANSP). ICES Scientific Reports. 10.17895/ices.pub27909783.v2

Karahan, A., Borsa, P., Gucu, A. C., Kandemir, I., Ozkan, E., Orek, Y. A., Acan, S. C., Koban, E., & Togan, I. (2014). Geometric morphometrics, Fourier analysis of otolith shape, and nuclear-DNA markers distinguish two anchovy species (Engraulis spp.) in the Eastern Mediterranean Sea. Fisheries Research, 159, 45–55. 10.1016/j.fishres.2014.05.009

Kerr, L. A., Hintzen, N. T., Cadrin, S. X., Clausen, L. W., Dickey-Collas, M., Goethel, D. R., Hatfield, E. M. C., Kritzer, J. P., & Nash, R. D. M. (2016). Lessons learned from practical approaches to reconcile mismatches between biological population structure and stock units of marine fish. ICES Journal of Marine Science, 74(6), 1708–1722 10.1093/icesjms/fsw188

Kerr, L. A., Hintzen, N. T., Cadrin, S. X., Clausen, L. W., Dickey-Collas, M., Goethel, D. R., Hatfield, E. M. C., Kritzer, J. P., & Nash, R. D. M. (2017). Lessons learned from practical approaches to reconcile mismatches between biological population structure and stock units of marine fish. ICES Journal of Marine Science, 74(6), 1708–1722. 10.1093/icesjms/fsw188

Le Moan, A., Gagnaire, P. A., & Bonhomme, F. (2016). Parallel genetic divergence among coastal-marine ecotype pairs of European anchovy explained by differential introgression after secondary contact. Mol Ecol, 25(13), 3187–3202. 10.1111/mec.13627

Leone, A., Álvarez, P., García, D., Saborido-Rey, F., & Rodriguez-Ezpeleta, N. (2019). Genome-wide SNP based population structure in European hake reveals the need for harmonizing biological and management units. ICES Journal of Marine Science 10.1093/icesjms/fsz161/5561460

Lischer, H. E., & Excoffier, L. (2012). PGDSpider: an automated data conversion tool for connecting population genetics and genomics programs. Bioinformatics, 28(2), 298–299. 10.1093/bioinformatics/btr642

Luu, K., Bazin, E., & Blum, M. G. B. (2016). Pcadapt: an R package to perform genome scans for selection based on principal component analysis. Molecular Ecology Resources, 17(1). 10.1101/056135

Magoulas, A., Castilho, R., Caetano, S., Marcato, S., & Patarnello, T. (2006). Mitochondrial DNA reveals a mosaic pattern of phylogeographical structure in Atlantic and Mediterranean populations of anchovy (Engraulis encrasicolus). Mol Phylogenet Evol, 39(3), 734–746. 10.1016/j.ympev.2006.01.016

Meyer, L., Barry, P., Le Moan, A., Arbiol, C., Castilho, R., Van der Lingen, C., Chlaïda, M., McKeown, N., Ernande, B., Bonhomme, F., Gagnaire, P. A., & Guinand, B. (2025). Genome divergence between European anchovy ecotypes fuelled by structural variants originating from trans-equatorial admixture. Proc Biol Sci, 292(2058), 20251416. 10.1098/rspb.2025.1416

Montes, I., Zarraonaindia, I., Iriondo, M., Grant, W. S., Manzano, C., Cotano, U., Conklin, D., Irigoien, X., & Estonba, A. (2016). Transcriptome analysis deciphers evolutionary mechanisms underlying genetic differentiation between coastal and offshore anchovy populations in the Bay of Biscay. Marine Biology, 163(10). 10.1007/s00227-016-2979-7

Oueslati, S., Fadhlaoui-Zid, K., Kada, O., Augé, M. T., Quignard, J. P., & Bonhomme, F. (2014). Existence of two widespread semi-isolated genetic entities within Mediterranean anchovies. Marine Biology, 161(5), 1063–1071. 10.1007/s00227-014-2399-5

Petitgas, P., Alheit, J., Peck, M. A., Raab, K., Irigoien, X., Huret, M., van der Kooij, J., Pohlmann, T., Wagner, C., Zarraonaindia, I., & Dickey-Collas, M. (2012). Anchovy population expansion in the North Sea. Marine Ecology Progress Series, 444, 1–13. 10.3354/meps09451

Pujolar, J. M., Gardiner, C. E. C., von der Heyden, S., Robalo, J. I., Castilho, R., Cunha, R., Henriques, R., & Nielsen, E. E. (2025). Resolving the Population Structure and Demographic History of the European Anchovy in the Northeast Atlantic: Tracking Historical and Contemporary Environmental Changes. Mol Ecol, e17829. 10.1111/mec.17829

Purcell, S., Neale, B., Todd-Brown, K., Thomas, L., Ferreira, M. A., Bender, D., Maller, J., Sklar, P., de Bakker, P. I., Daly, M. J., & Sham, P. C. (2007). PLINK: a tool set for whole-genome association and population-based linkage analyses. Am J Hum Genet, 81(3), 559–575. 10.1086/519795

Reid, J. L. (1966). Oceanic environments of the genus Engraulis around the world. CalCOFI, Rep. 11, 29–33.

Reiss, H., Hoarau, G., Dickey-Collas, M., & Wolff, W. J. (2009). Genetic population structure of marine fish: mismatch between biological and fisheries management units. Fish and Fisheries, 10(4), 361–395. 10.1111/j.1467-2979.2008.00324.x

Ruiz, J., González-Quirós, R., Prieto, L., & Navarro, G. (2009). A Bayesian model for anchovy (Engraulis encrasicolus): the combined forcing of man and environment. Fisheries Oceanography, 18(1), 62–76. 10.1111/j.1365-2419.2008.00497.x

Sanz, N., García-Marín, J.-L., Viñas, J., Roldán, M., & Pla, C. (2008). Spawning groups of European anchovy: population structure and management implications. ICES Journal of Marine Science, 65(9), 1635–1644. 10.1093/icesjms/fsn128

Silva, G., Horne, J. B., Castilho, R., & Maggs, C. (2014). Anchovies go north and west without losing diversity: post-glacial range expansions in a small pelagic fish. Journal of Biogeography, 41(6), 1171–1182. 10.1111/jbi.12275

Silva, G., Lima, F. P., Martel, P., & Castilho, R. (2014). Thermal adaptation and clinal mitochondrial DNA variation of European anchovy. Proc Biol Sci, 281(1792). 10.1098/rspb.2014.1093

Teles-Machado, A., Plecha, S. M., Peliz, A., & Garrido, S. (2024). Anomalous ocean currents and European anchovy dispersal in the Iberian ecosystem. Marine Ecology Progress Series, 741, 289–300. 10.3354/meps14526

Therkildsen, N. O., & Palumbi, S. R. (2017). Practical low-coverage genomewide sequencing of hundreds of individually barcoded samples for population and evolutionary genomics in nonmodel species. Molecular Ecology Resources, 17(2), 194–208. 10.1111/1755-0998.12593

van der Kooij, J., McKeown, N., Campanella, F., Boyra, G., Doray, M., Santos Mocoroa, M., Fernandes da Silva, J., & Huret, M. (2024). Northward range expansion of Bay of Biscay anchovy into the English Channel. Marine Ecology Progress Series, 741, 217–236. 10.3354/meps14603

Viñas, J., Sanz, N., Peñarrubia, L., Araguas, R.-M., García-Marín, J.-L., Roldán, M.-I., & Pla, C. (2014). Genetic population structure of European anchovy in the Mediterranean Sea and the Northeast Atlantic Ocean using sequence analysis of the mitochondrial DNA control region. ICES Journal of Marine Science, 71(2), 391–397. 10.1093/icesjms/fst132

Zarraonaindia, I., Iriondo, M., Albaina, A., Pardo, M. A., Manzano, C., Grant, W. S., Irigoien, X., & Estonba, A. (2012). Multiple SNP markers reveal fine-scale population and deep phylogeographic structure in European anchovy (Engraulis encrasicolus L.). Plos One, 7(7), e42201. 10.1371/journal.pone.0042201

Zarraonaindia, I., Pardo, M. A., Iriondo, M., Manzano, C., & Estonba, A. (2009). Microsatellite variability in European anchovy (Engraulis encrasicolus) calls for further investigation of its genetic structure and biogeography. ICES Journal of Marine Science, 66(10), 2176–2182. 10.1093/icesjms/fsp187

